# Time-Resolved Laser Speckle Contrast Imaging (TR-LSCI) of Cerebral Blood Flow Response to Intracranial Pressure Elevation

**DOI:** 10.64898/2026.02.19.706870

**Authors:** Faraneh Fathi, Peiwen Zhang, Mehrana Mohtasebi, Paul Mos, Claudio Bruschini, Edoardo Charbon, Jin Chen, Li Chen, Guoqiang Yu, Lei Chen

**Author notes:** Corresponding Authors., Guoqiang Yu, Lei Chen.

## Abstract

**Significance:** Cerebral autoregulation (CA) reflects the dynamic coupling among cerebral blood flow (CBF), intracranial pressure (ICP), and arterial blood pressure (ABP); its failure contributes to secondary brain injury. Existing bedside methods rely on indirect or spatially limited CBF surrogates and cannot resolve microvascular flow dynamics across space, depth, and time.

**Aim:** To develop, optimize, and apply a scalable, noncontact time-resolved laser speckle contrast imaging (TR-LSCI) platform for depth-sensitive, high-speed, wide-field CBF imaging during controlled ICP perturbations.

**Approach:** TR-LSCI synchronizes a 20-MHz pulsed laser with a time-gated, single-photon avalanche diode (SPAD) camera (512 × 512 pixels) to detect diffuse photons at varying path lengths, enabling depth-resolved microvascular CBF imaging. Benchtop and mobile TR-LSCI systems were applied in adult rats and a neonatal piglet with synchronized invasive ICP and ABP measurements.

**Results:** TR-LSCI captured spatially heterogeneous, pulsatile CBF dynamics at up to 52 Hz over large cortical fields of view, with heart rate estimates statistically equivalent to those from ICP and ABP. Multivariable analysis identified reproducible, phase-dependent CA transitions encompassing preserved autoregulation, ABP-driven compensation, and ICP-constrained CBF suppression; notably, CBF alone exhibited distinct phase signatures.

**Conclusions:** TR-LSCI enables dynamic, physiology-informed neurovascular monitoring and supports future bedside CA assessment.

## 1 Introduction

Cerebral blood flow (CBF) is the primary determinant of cerebral oxygen delivery and metabolic viability, and its regulation is critically influenced by intracranial pressure (ICP) and arterial blood pressure (ABP), together forming a tightly coupled physiological triad. Disruption of this triad plays a central role in secondary brain injury in a wide range of neuropathologies, including traumatic brain injury, intracranial hemorrhage, hydrocephalus, and ischemic stroke [1–10]. Elevated or unstable ICP compromises cerebral perfusion pressure (CPP), alters vascular compliance, and is strongly associated with poor neurological outcomes [3, 8]. Consequently, understanding how CBF responds to changes in ICP and ABP under dynamic pathological conditions is fundamental to neurocritical care.

The physiological relationship among CBF, ICP, and ABP is grounded in the Monro-Kellie doctrine, which posits that the cranial vault is a fixed-volume compartment consisting of brain tissue, blood, and cerebrospinal fluid [11, 12]. Changes in any one component must be offset by others, leading to altered intracranial compliance, vascular transmural pressure, and cerebral perfusion. Together, these relationships formalize how ICP and ABP jointly govern CBF. Both experimental and mathematical models have demonstrated that progressive ICP elevation reshapes cerebrovascular impedance and alters the transmission of cardiac pulsatility through the cerebral vasculature, thereby modulating both ICP and CBF waveforms [13, 14]. These coupled mechanisms suggest that CBF dynamics encode rich information about the evolving intracranial mechanical and hemodynamic state, beyond static perfusion levels alone.

Within this framework, cerebral autoregulation (CA) represents the principal protective mechanism governing this triad, enabling cerebral arterioles to adjust vascular tone and maintain relatively stable CBF despite fluctuations in CPP [1, 15, 16]. Classical descriptions of CA, most notably the Lassen curve, proposed a broad plateau over which CBF remains pressure-independent and approximately constant across an ABP range of ∼50-150 mmHg [17, 18]. However, clinical and experimental evidence now indicates that the effective autoregulatory range is substantially narrower than originally proposed and varies across disease states, developmental stages, and physiological conditions such that even modest perturbations in ABP or ICP can measurably alter CBF [19]. These observations have shifted contemporary views of CA toward a dynamic, multivariate, and time-dependent process governed by interacting myogenic, neurogenic, endothelial, and metabolic mechanisms rather than ABP alone [16, 20–22].

This evolving understanding has motivated growing interest in continuous bedside monitoring of CA to guide individualized ABP management in neurocritical care. However, fixed, population-level ABP targets recommended by current guidelines may fail to account for inter-patient variability in cerebrovascular reserve and may predispose vulnerable patients to hypoperfusion or hyperemia when autoregulation is impaired [2, 23, 24]. As emphasized in recent consensus efforts, real-time assessment of cerebrovascular state is increasingly viewed as essential for personalized perfusion management [16].

Modern dynamic CA monitoring approaches estimate autoregulatory function by correlating spontaneous low-frequency fluctuations in ABP or CPP with surrogate measures of cerebral hemodynamics. Commonly used indices include the pressure reactivity index (PRx), derived from correlations between ICP and ABP as a proxy for cerebral blood volume reactivity [25–27]. Transcranial Doppler (TCD) based indices such as mean flow index (Mx), which correlate large vessel CBF velocity with ABP or CPP [28–31] and near-infrared spectroscopy (NIRS) based indices such as cerebral oximetry index (Cox), tissue oxygenation index (Tox), tissue hemoglobin index (THx), and hemoglobin volume index (HVx), which relate cerebral oxygenation or hemoglobin signals to perfusion pressure [32–35]. Diffuse correlation spectroscopy (DCS) has further enabled frequency-domain and transfer-function based CA metrics, including gain and phase relationships between ABP and microvascular CBF, providing sensitivity to both the magnitude and latency of autoregulatory responses [36–40].

Despite these advances, all existing bedside CA monitoring modalities rely on indirect or spatially restricted surrogates of CBF. PRx and related ICP-based indices require invasive electrode monitoring and provide global rather than regional information in the brain [41, 42]. TCD measures flow velocity in large intracranial arteries. However, its clinical utility is limited by reliance on large vessels, operator dependence, challenges in longitudinal monitoring, and poor acoustic windows, particularly in neonates [43, 44]. NIRS and DCS directly probe cerebral oxygenation and CBF and enable dynamic CA analysis, but they remain inherently contact-dependent with limited spatiotemporal resolution, restricting their ability to resolve rapid and heterogeneous cortical responses [45–48]. Accordingly, existing noninvasive modalities do not fully provide continuous, noncontact, depth-resolved microvascular CBF measurements with both high spatial and temporal resolution over large head coverage, limiting their suitability for comprehensive dynamic assessment of CA.

To address this gap, we recently developed a time-resolved laser speckle contrast imaging (TR-LSCI) technology that synchronizes pulsed, wide-field laser illumination with a gated single-photon avalanche diode (SPAD) camera, enabling noncontact, depth-resolved, high-speed, and high-density CBF imaging over a large head area. The first benchtop TR-LSCI prototype demonstrated depth-resolved CBF imaging in head-mimicking phantoms and rodent models [49]. The system was subsequently optimized by upgrading the gated camera from SwissSPAD2 to SPAD512^2^ and integrating a latent diffusion model to suppress diffusive noise [50]. Most recently, the TR-LSCI platform was redesigned as a cart-mounted, mobile device with fiber-coupled illumination and a compact, rotatable probe head to facilitate clinical translation [51].

In this study, we first conducted adult rat experiments using the benchtop TR-LSCI system to establish feasibility and characterize CBF responses to ICP and ABP perturbations under tightly controlled conditions. Specifically, a continuous, graded intraventricular saline infusion protocol was used to induce a wide range of ICP variations while acquiring high-speed measurements of CBF, ICP, and ABP using noninvasive TR-LSCI and invasive pressure sensors, respectively. To enhance translational relevance, we subsequently extended the study to a neonatal piglet model using the mobile TR-LSCI platform. Unlike the rat protocol, piglet experiments employed stepwise, non-continuous saline injections with defined infusion and pause recovery periods, allowing ICP to rise and stabilize between steps and enabling assessment of both transient and sustained CA responses under clinically realistic conditions.

Results from this study demonstrate that TR-LSCI enables noncontact, high-speed (up to 52 Hz in the present study), high-density (up to 512 × 512 pixels), and depth-sensitive imaging of CBF variations across a large cortical field of view (FOV). This capability enables direct visualization of spatially heterogeneous CBF responses over time during dynamic ICP and ABP perturbations, which is not achievable with existing technologies. By concurrently monitoring CBF, ICP, and ABP, we show that cerebrovascular regulation evolves through multiple distinct physiological phases, beginning with intact CA and followed by a transition from ABP-driven CBF augmentation to ICP-driven CBF suppression. These findings establish a new experimental framework for elucidating the dynamic coupling among ICP, ABP, and CBF, providing a foundation for developing noninvasive optical strategies for continuous CA monitoring.

## 2 Methods and Materials

### 2.1 In Vivo Experimental Protocols

All experimental procedures were approved by the Institutional Animal Care and Use Committee (IACUC) at the University of Kentucky. Two complementary in vivo studies were conducted to evaluate TR-LSCI performance and cerebrovascular regulation under controlled ICP perturbations: (I) a rodent model enabling continuous graded ICP loading and (II) a neonatal piglet model designed to assess translational feasibility in a large-animal setting. **Fig 1** shows the experimental configuration and instrumentation for both studies.

**Fig. 1.**
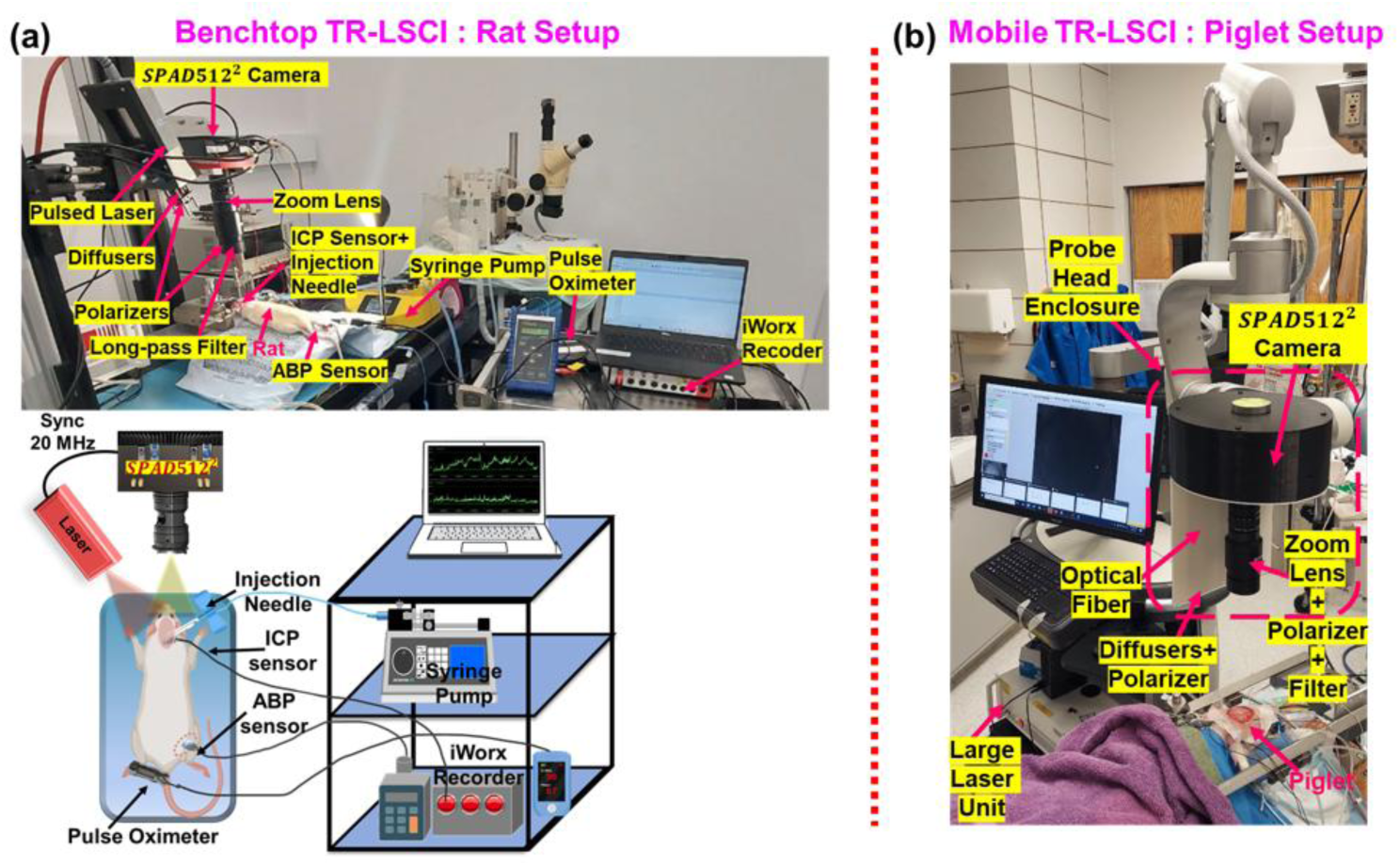
TR-LSCI system configuration and portability optimization for imaging pulsatile CBF waves in adult rats and a neonatal piglet. **(a)** Benchtop TR-LSCI system for CBF imaging in adult rats. The setup shows an open-space pulsed laser with diffusers synchronized with the SPAD512² camera, zoom lens, long-pass filter, and polarization optics. The schematic also illustrates integration of multimodal physiological measurements, including ICP and ABP sensors, a pulse oximeter, and data acquisition units. A syringe pump was used to deliver controlled intraventricular saline infusion for graded ICP modulation. (b) Mobile fiber-coupled TR-LSCI system for CBF imaging in a neonatal piglet. The portable probe head housed the SPAD512² camera within a custom 3D-printed enclosure, with the zoom lens, polarizer, and long-pass filter mounted externally. The pulsed laser light was coupled to and delivered through a 6-m multimode optical fiber, enabling full separation of the large laser unit from the portable probe head. The articulated support arm and probe enclosure enabled rapid, stable positioning while preserving precise optical alignment for translational measurements.

#### Rat Study (Fig. 1a)

Four adult male Sprague-Dawley rats (2-3 months old) were used to characterize cerebrovascular responses to controlled ICP modulation. Animals were anesthetized with 1-2% isoflurane, placed on a thermostatically controlled heating blanket, and secured in a stereotaxic frame. Peripheral blood oxygen saturation (SpO_2_) and heart rate (HR) were continuously monitored using a pulse oximeter (mouseO_x_ pluse, STARR Life Sciences Corp), and body temperature was maintained within the normal range.

A commercial fiber-optic pressure sensor (OPsens, Inc.) was inserted into the left femoral artery for continuous ABP monitoring. The scalp was removed to expose the intact skull, which was sealed with a thin layer of mineral oil to maintain tissue hydration. A commercial solid-state ICP sensor (A-BP-CATH-16, 1.6 F, 0.53 mm outer diameter, iWorx Systems, Inc.) and a 22G injection needle were inserted into the right lateral ventricle through a small burr hole in the right temporal skull. ICP values were continuously recorded using a multichannel data acquisition system (IX-RA-834 10+ Channel Recorder and Stimulator, iWorx Systems, Inc.). After a 15-min stabilization period, noncontact TR-LSCI imaging was initiated and continued throughout controlled ICP modulation. TR-LSCI was performed over a cortical region of approximately 34 × 17 mm^2^, enabling continuous imaging of blood flow index (BFI) and extraction of relative CBF (rCBF) changes.

Following a 5 to 7 min baseline measurement, saline was infused continuously through the ventricular needle in a graded sequence. The initial infusion rate of 0.05 mL/min was maintained for 20 min. Infusion rates were then increased to 0.1, 0.2, and 0.3 mL/min, each maintained for 5 min. The 1.0 mL/min step was applied with duration determined by physiological tolerance. In one animal (Rat #2), the 1.0 mL/min infusion was limited to 3 min and escalation to 3.0 mL/min was not performed. In the remaining animals, a final high-rate infusion (3.0 mL/min) was continued until terminal conditions were reached, with duration varying across animals depending on physiological tolerance. In one of the earliest animals studied, the 0.05 mL/min step was not applied. This protocol enabled systematic interrogation of dynamic coupling among ICP, ABP, and CBF under controlled physiological perturbations.

#### Piglet Study (Fig. 1b)

To enhance translational relevance, experiments were extended to a large animal model of neonatal piglets. One male neonatal piglet (postnatal day 7; genetic background of mixed White Yorkshire and Landrace) was obtained from Oak Hill Genetics through the Division of Laboratory Animal Resources (DLAR) at the University of Kentucky. Animals were housed in a temperature-controlled enclosure and fed ad libitum with milk replacer (Birthright, Ralco), with fasting for at least 4 hours prior to surgery.

Sedation was induced with Midazolam (0.1-0.5 mg/kg), followed by intubation using a pediatric endotracheal tube (3-mm internal diameter) and maintenance under 1.5-2% isoflurane anesthesia. The piglet was positioned in a customized stereotaxic frame and maintained at normothermia using a circulating water pad and Bair Hugger system, with continuous temperature monitoring via a rectal probe. A multichannel respirator-oximeter system (8400, Smiths Medical) recorded HR, respiration rate, and SpO_2_ throughout the experiment.

A solid-state pressure sensor (A-BP-CATH-16, 1.6 F, 0.53 mm outer diameter, iWorx Systems, Inc.) was surgically inserted into the left femoral artery for ABP monitoring. After surgical preparation, the scalp was removed to expose the skull, and mineral oil was applied to maintain tissue hydration. A small burr hole (∼2 mm) was drilled in the right temporal bone, through which a solid-state ICP probe (A-BP-CATH-48, 4.8 F, 1.58 mm outer diameter, iWorx Systems, Inc.) and a 22G blunt-tip needle were inserted into the right lateral ventricle. After a 15-minute stabilization period, noncontact TR-LSCI imaging was continuously performed over an intact-skull region of approximately 47 × 47 mm² with simultaneous ICP and ABP acquisition.

The piglet intracranial loading protocol consisted of phasic saline infusions with defined injection and pause periods. Following a baseline recording, saline was infused at 0.2 mL/min until ICP reached 30 mmHg (∼10 minutes). The infusion was then stopped for 14 minutes to allow ICP to return to baseline. A second infusion was administered at 0.2 mL/min and controlled to reach an ICP target of 30 mmHg (∼5 minutes). This was followed by a 5-minute pause, after which the infusion rate was increased to 0.5 mL/min for 6 minutes. The experiment concluded with a terminal phase and euthanasia with pentobarbital overdose over 2 minutes. This protocol produced both transient and sustained intracranial pressure elevations under clinically relevant conditions and enabled direct comparison of cerebrovascular regulation between the rodent and large-animal neonatal models.

### 2.2 TR-LSCI Systems and Portability Optimization

**Fig. 1** shows the benchtop TR-LSCI system configuration for adult rat experiments and the corresponding mobile, fiber-coupled TR-LSCI platform developed for neonatal piglet imaging, including the optical layout, synchronized physiological monitoring, and infusion control.

#### Benchtop TR-LSCI System (Fig 1a)

The benchtop TR-LSCI system used in this study for the rat experiments was adapted from our previously reported benchtop platform for noncontact, depth-resolved, wide-field CBF imaging [50]. The system employed a time-gated SPAD512² camera (Pi Imaging, Switzerland) featuring a 512 × 512 pixel sensor array. Coherent pulsed illumination at 20 MHz was provided by a pulsed 775 nm laser (Katana-08 HP, NKT Photonics), diffused using two engineered diffusers (ED1-C20-MD and ED1-S20-MD, Thorlabs) to achieve spatially uniform illumination. A Zoom 7000 lens (Navitar) provided flexible region-of-interest (ROI) control, while polarization optics in both the illumination and detection paths suppressed surface glare. A >750 nm long-pass filter (84-761, Edmund Optics) reduced ambient light contamination.

#### Mobile TR-LSCI Device (Fig. 1b)

To enable large-animal imaging and translational studies, the TR-LSCI system was subsequently redesigned as a mobile, fiber-coupled imaging platform for piglet experiments [51]. A custom 3D-printed probe enclosure was fabricated to house the SPAD512² camera, while the zoom lens, polarizer, and long-pass filter were mounted externally and extended from the enclosure, allowing flexible optical and mechanical adjustment and simplified maintenance. The pulsed laser source was coupled into a multimode optical fiber (200-µm core, 0.39 NA; Thorlabs FT200UMT) via an SMA coupler (F110SMA-780, Thorlabs), with an approximately 6-m fiber length, enabling complete separation of the large laser unit from the portable probe head. The same diffuser, polarization, and filter optics used in the benchtop configuration were integrated into the portable probe head.

In the portable probe design, the fiber-coupled output was positioned laterally and as close as possible to the base of the camera zoom lens to minimize the side-illumination angle relative to the imaging axis while reducing illumination shadowing. The articulated support arm and probe enclosure provided multiple degrees of freedom, enabling rapid and stable positioning over the target while maintaining consistent optical alignment.

#### TR-LSCI System Configuration and Data Acquisition

System configuration was guided by our prior TR-LSCI studies in head-mimicking phantoms and rodent brain experiments with well-characterized optical and geometric parameters [49–51]. Briefly, the TR-LSCI system operated in gated mode with laser-camera synchronization at 20 MHz (50 ns period). Depth-resolved sampling was achieved by shifting the detection gate with a minimal delay of 18.6 ps. The gate width was fixed at 13 ns for both rat and piglet experiments.

For rat experiments, 8-bit frames were acquired with 1.3 ms exposure. The raw image size was configured to 512 × 256 pixels to increase frame rate while maintaining sufficient cortical coverage. To preserve high temporal resolution for capturing cardiac pulsatility, the number of depth gates was limited to three, since additional gates would reduce the effective sampling rate and degrade waveform fidelity. With an experimentally determined initial gate offset of 44,740 ps, three gates (Gate #0, #35, and #55) were generated using delay increments of 0, 35 × 18.6 ps, and 55 × 18.6 ps, corresponding to separations of 0, ∼0.65 ns, and ∼1.02 ns. This configuration balanced cortical sensitivity and depth separation through the thin adult rat skull while maintaining a frame rate of ∼52 Hz. rCBF was derived from Gate #0, which provided sufficient coverage of cortical tissue under the thin skull of rats.

For piglet experiments, 8-bit frames were acquired with 1.8 ms exposure using the full sensor size of 512 × 512 pixels to maximize FOV over the larger cranial surface. An initial gate offset of 48,267 ps and three depth gates (**Gate #0, #40, and #65**) were applied, with larger delay increments of 0, 40 × 18.6 ps, and 65 × 18.6 ps, corresponding to separations of 0, ∼0.74 ns and ∼1.21 ns. These settings shifted the detection window later in time to improve penetration depth and signal-to-noise ratio (SNR) under the thicker skull and larger head geometry of the piglet, while preserving a sampling rate of ∼46 Hz. rCBF was primarily derived from Gate #40, selected based on prior TR-LSCI depth-sensitivity characterization in head-mimicking phantoms and the 1-2 mm skull thickness reported in preterm and term neonatal piglets [49, 51, 52]. Gate-dependent analysis further confirmed higher absolute BFI values and stronger deep-tissue contributions at later gates, consistent with increased photon pathlength and depth sensitivity [51]. Overall, these acquisition strategies enabled depth-aware, high-speed BFI imaging across both animal models while preserving robust pulsatile rCBF signal fidelity.

### 2.3 Signal Processing and Data Analysis Pipeline

**Fig. 2** summarizes the complete signal-processing and data analysis pipeline used in this study, encompassing **(a)** conversion of depth-gated TR-LSCI intensity images into spatial BFI maps and ROI-based rCBF time courses, **(b)** synchronized acquisition and segmentation of ICP and ABP signals using experimental markers, **(c)** baseline cardiac pulsatility analysis and heart-rate extraction from rCBF, ICP, and ABP, and **(d)** BFI, ICP and ABP interaction analysis during graded intracranial infusion.

**Fig. 2.**
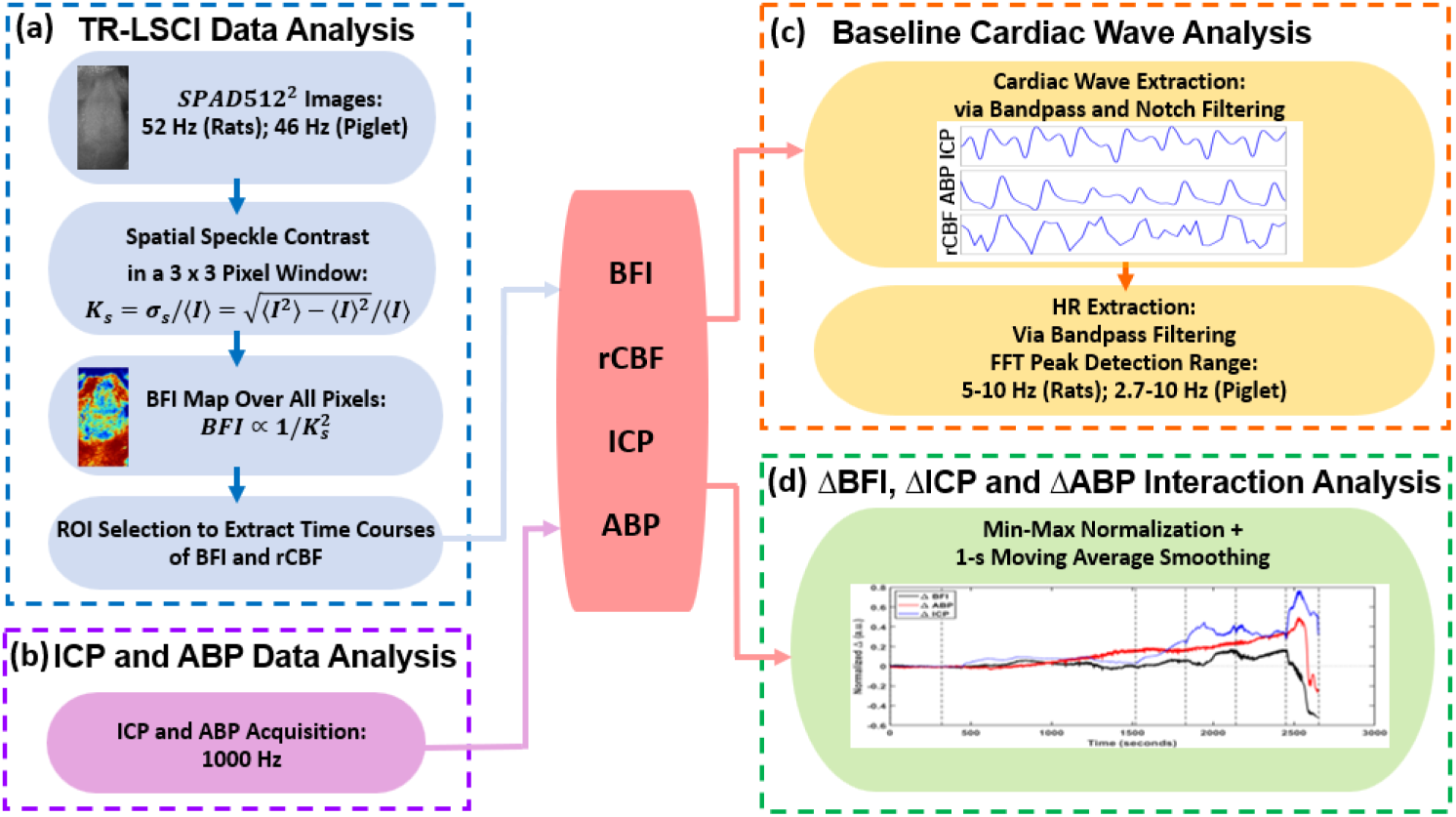
Flowchart summarizing TR-LSCI and multimodal physiological data-analysis pipeline. **(a)** Raw gated SPAD512² intensity images acquired by TR-LSCI (∼52 Hz in rats and ∼46 Hz in piglets) are converted to spatial speckle-contrast maps using a sliding 3×3 pixel window, from which two-dimensional BFI maps are estimated and ROI-averaged rCBF time courses are extracted and normalized to the first 100 s of baseline. **(b)** ICP and ABP data acquired at 1000 Hz and aligned with BFI based on experiment markers. **(c)** Baseline cardiac pulsatile dynamics analysis using species-specific bandpass filtering, followed by FFT-based heart-rate extraction using sliding windows. **(d)** BFI, ICP, and ABP interaction analysis. To examine pressure-flow interactions during graded intracranial loading, BFI, ICP, and ABP are min-max normalized and converted to baseline-relative form, enabling joint visualization of low-frequency ΔBFI, ΔICP, and ΔABP trends and identification of phase-dependent cerebrovascular regulatory triad.

#### TR-LSCI Data Analysis (Fig. 2a)

TR-LSCI data processing followed our previously described pipeline [34,35]. Raw SPAD512^2^ intensity frames were converted to speckle-contrast maps using spatial LSCI analysis. For each pixel, the spatial speckle contrast 𝐾_𝑠_ was computed in a sliding 3 × 3 window as 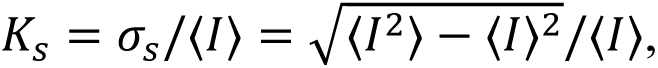 where 𝜎_𝑠_ and ⟨𝐼⟩ denote the local standard deviation and mean intensity, respectively. A blood flow index (BFI) was then estimated using the established inverse-square relationship BFI ∝ 1/𝐾_s_^2^. A two-dimensional BFI map was generated over all pixels. To improve SNR, BFI maps over the final 20 s of each measurement segment were averaged for visualization of spatial CBF distribution. A cortical ROI was selected on the BFI maps, and the ROI-averaged BFI value from each frame was extracted to form a continuous time series. This series was normalized to the mean BFI during the first 100 s of baseline to obtain rCBF, enabling analysis of pulsatile rCBF dynamics. The effective rCBF sampling frequency was determined by the acquisition configuration and differed by animal models: ∼52 Hz in rats and ∼46 Hz in the piglet.

#### ICP and ABP Data Analysis (Fig. 2b)

ICP and ABP were acquired continuously at 1000 Hz using solid-state and fiber-optic pressure sensors and synchronized with TR-LSCI via experiment markers defining baseline and infusion epochs. Because each modality was analyzed and visualized at its native sampling rate, no data interpolation was required.

#### Baseline Cardiac Pulsatile Wave Analysis (Fig. 2c)

Following extraction of full-length time-course data, baseline cardiac pulsatile dynamics were examined to characterize intrinsic cardiac pulsatility under stable conditions. For adult rat recordings, a bandpass filter was applied to rCBF, ICP, and ABP signals to suppress low-frequency respiratory oscillations while preserving cardiac pulsatility and its harmonics. Specifically, a fourth-order zero-phase infinite impulse response (IIR) bandpass filter with half-power cutoffs at 3 Hz and 25 Hz was applied, effectively removing respiratory components (<3 Hz) while retaining cardiac-frequency content (≥5 Hz) and higher-order harmonics relevant for pulsatile waveform analysis. HR was then extracted from rCBF, ICP, and ABP, respectively, using an overlapping-window FFT approach. Signals were analyzed in 30-s windows with 0.1-s hops, and HR was extracted as the dominant spectral peak within the 5-10 Hz band, consistent with rat heart physiology. HR estimates were computed independently for each modality at their native sampling rates and then temporally aligned for joint statistical analysis.

For neonatal piglet recordings, ICP signals contained narrowband 60 Hz electrical line noise, which was removed using a fourth-order IIR notch filter (50-70 Hz). ICP and ABP were then bandpass filtered (fourth-order, zero-phase; 2.5-25 Hz) to extract HR using an overlapping-window FFT. rCBF signals, which showed no 60 Hz contamination, were filtered using the same bandpass strategy at their native sampling rate (46 Hz), with an upper cutoff of 23 Hz to satisfy the Nyquist criterion. HR was extracted as the dominant spectral peak within the 2.7-10 Hz band, consistent with neonatal piglet cardiac physiology. Occasional outliers in ICP-derived HR estimates were removed using a robust median absolute deviation (MAD) criterion applied jointly across ABP, ICP, and rCBF; values exceeding three MADs from the median were excluded to ensure physiologically consistent baseline HR measurements.

#### BFI, ICP and ABP Interaction Analysis (Fig. 2d)

To examine interactions among BFI, ICP, and ABP during graded intracranial pressure loading, these three signals were analyzed as comparable slow-trend trajectories across infusion stages. For joint visualization, each signal was min-max normalized over the analysis interval and converted to baseline-relative form by subtracting the mean value of the first 100 s at the baseline. This transformation removes differences in physical units and dynamic range while highlighting deviations (Δ) from baseline.

A one-second moving average was optionally applied to reduce residual high-frequency fluctuations and improve readability, while preserving physiologically meaningful slow dynamics. These signals were then plotted together to enable qualitative comparison of ΔBFI and CA variations relative to concurrent ΔABP and ΔICP fluctuations across infusion stages.

### 2.4 Statistical Analysis

All statistical analyses were performed in MATLAB. Equivalence testing, rather than traditional difference testing, was used to assess whether HR estimates derived from ABP, ICP, and rCBF signals were physiologically equivalent. Equivalence was evaluated using the two one-sided tests (TOST) procedure, implemented as two one-sided one-sample t-tests on the paired difference with a priori equivalence margin of ±0.25 Hz, defined as the largest physiologically negligible difference relative to typical animal HR variability and system resolution. A 90% confidence interval (CI) and significance level of α = 0.05 were used throughout, and equivalence was concluded only when both one-sided tests were significant and the 90% CI lay entirely within the equivalence bounds.

#### Across-Subject Analysis

For each rat (n = 4), mean baseline HR values across baseline time points were computed, yielding one averaged HR per modality per rat. Equivalence in averaged HR for ABP vs rCBF, ICP vs rCBF, and ABP vs ICP was evaluated using the TOST procedure as described above. An across-subject analysis was not feasible for the piglet cohort due to a single subject.

#### Within-Subject (Across-Time-Point) Analysis

Equivalence testing was also performed across baseline time points within each rat and the piglet. For each subject (rat/piglet), HR values at different baseline time points can be treated as independent paired observations, and the same three modality comparisons across time-points were evaluated using the TOST procedure as described above. In total, 15 equivalence tests were conducted (three comparisons per subject across four rats and one piglet), enabling assessment of equivalence robustness over time within individual subjects.

## 3 Results

This section presents experimental results obtained with TR-LSCI systems to characterize cerebrovascular responses to controlled ICP perturbations in adult rats and a neonatal piglet. First, rCBF imaging results acquired with the bench-top TR-LSCI system and synchronized physiological responses in adult rats are reported (**Section 3.1**), followed by translational results obtained in a larger neonatal piglet using the mobile TR-LSCI device (**Section 3.2**). Baseline cardiac pulsatility and heart-rate consistency across rCBF, ICP, and ABP measurements are then evaluated (**Section 3.3**). Finally, it is shown that combined monitoring of CBF, ICP, and ABP reveals cerebrovascular regulation evolving through multiple distinct CA phases during graded intraventricular infusion (**Section 3.4**).

### 3.1 rCBF, ABP, and ICP Dynamics in Adult Rats During Intraventricular Infusion

Representative TR-LSCI results from two adult rats are shown in **Fig. 3** and **Fig. 4**, illustrating rCBF responses (Gate #0) during graded intracranial pressure elevation. Each figure includes a raw TR-LSCI intensity image with the selected ROI, two-dimensional BFI maps, synchronized rCBF, ICP, and ABP time courses, and zoom-in pulsatile waveforms extracted from the baseline period of 1 second.

**Fig. 3.**
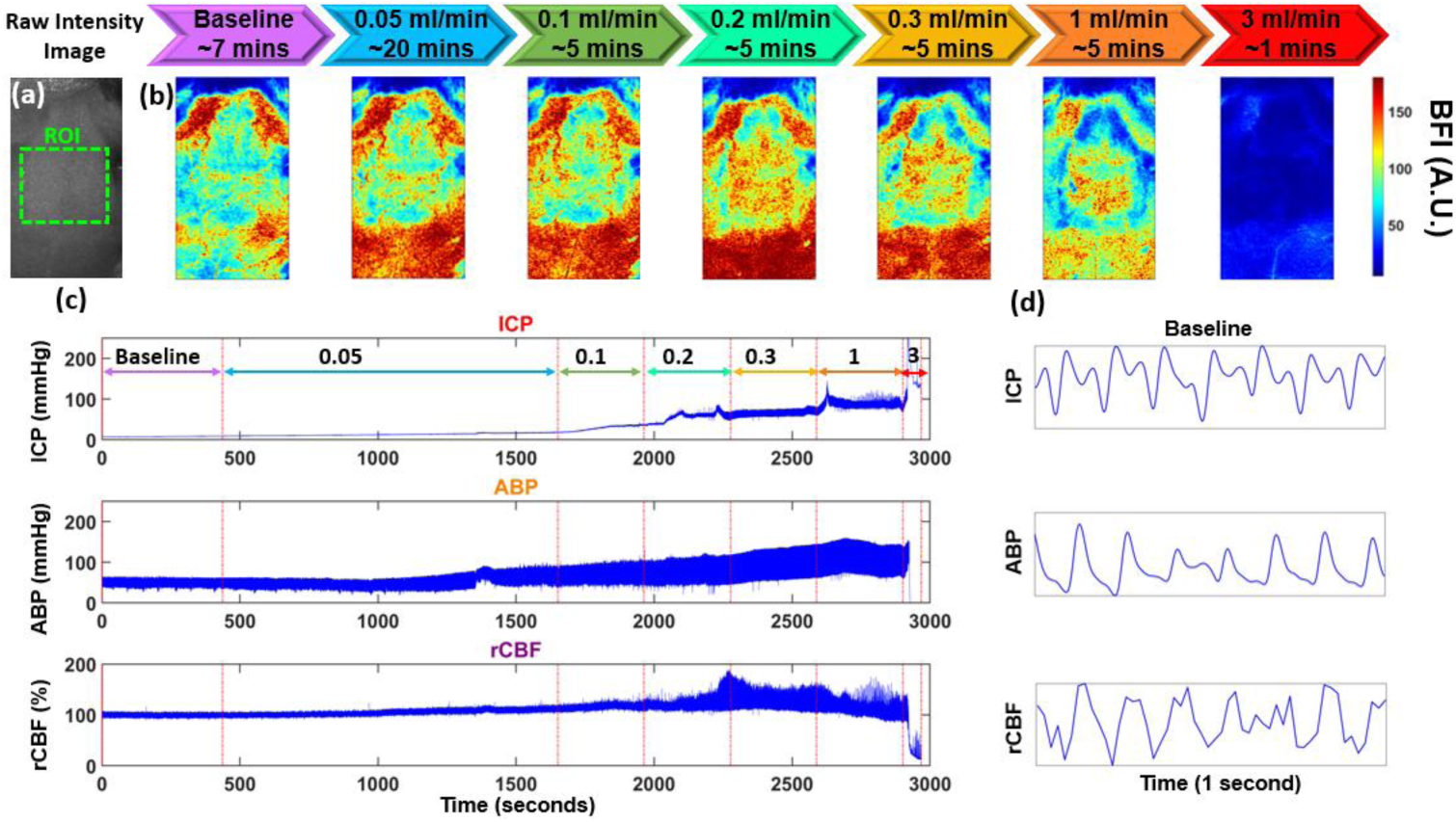
Representative rat results (Rat #1) demonstrating two-dimensional BFI maps and synchronized multimodal physiological responses during graded intracranial infusion. (a) Raw intensity image with the cortical ROI (green dashed box). **(b)** BFI maps acquired at baseline and during progressively increasing saline infusion rates of 0.05 ml/min (20 min), 0.1 ml/min (5 min), 0.2 ml/min (5 min), 0.3 ml/min (5 min), and 1.0 ml/min (5 min), and 3 ml/min (1min) illustrating spatial redistribution of cortical CBF with rising ICP. All BFI maps were derived from Gate #0 and represented averaging over the final 20 s of each infusion segment. **(c)** Corresponding time courses of ICP, ABP, and rCBF. Vertical dashed lines indicate infusion transitions. Sampling rates: ICP and ABP at 1000 Hz; rCBF at 52 Hz with three depth gates and 8-bit acquisition. **(d)** Zoom-in pulsatile waveforms of ICP, ABP, and rCBF extracted from the baseline segment of 1 sec following bandpass filtering.

**Fig. 4.**
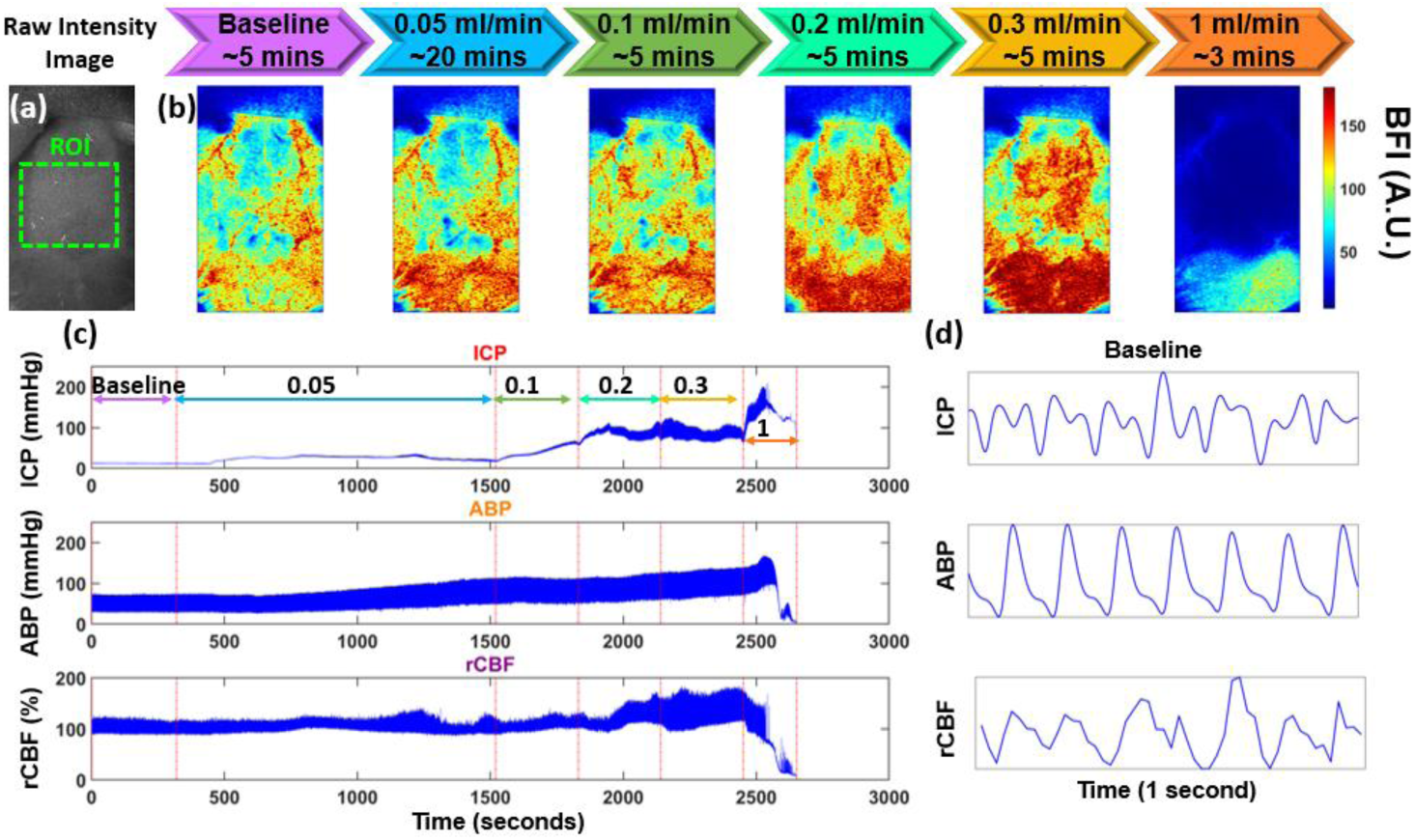
Reproducible pressure-flow dynamics in another representative rat (Rat #2) during graded intracranial infusion. **(a)** Raw intensity image with the selected ROI (green dashed box) for time-course rCBF analysis. **(b)** Two-dimensional BFI maps derived from Gate #0 during baseline and successive infusions. **(c)** Synchronized time courses of ICP, ABP, and rCBF across the graded infusion protocol. Vertical dashed lines indicate infusion transitions. Sampling rates: ICP and ABP at 1000 Hz; rCBF at 52 Hz with three depth gates and 8-bit acquisition. **(d)** Zoom-in pulsatile waveforms of ICP, ABP, and rCBF extracted from the baseline segment of 1 sec following bandpass filtering.

**Fig. 3** presents the results from a representative adult rat (Rat #1) during intracranial infusion. A cortical ROI was defined and used consistently for rCBF time-course extraction (**Fig. 3a**). The lower FOV primarily includes extracranial muscle, whereas the selected ROI encompasses exposed cortex; accordingly, all reported rCBF analyses are restricted to this cortical region.

At baseline, BFI exhibited a relatively spatially uniform distribution across the exposed skull (**Fig. 3b**). Following initiation of saline infusion into the right lateral ventricle, cortical BFI progressively increased and became spatially heterogeneous in a manner strongly dependent on infusion rate. During the prolonged low-rate infusion (0.05 ml/min for 20 min), cortical BFI remained stable or increased gradually and relatively uniformly, indicating that preserved CA was gradually impaired under modest ICP elevations. As infusion rates increased to 0.1, 0.2, and 0.3 ml/min (each maintained for ∼5 min), cortical BFI rose further and exhibited increased spatial variability, with regions of elevated flow emerging predominantly in central and posterior cortical areas. At higher infusion rates (1.0 mL/min for approximately 5 min), cortical BFI showed redistribution and partial attenuation. During the final high-rate infusion (3.0 mL/min), which was continued only until terminal physiological instability, BFI demonstrated pronounced regional suppression, consistent with severe impairment of autoregulatory capacity.

The corresponding multimodal time courses demonstrate tight coupling between the infusion protocol and physiological responses (**Fig. 3c**). ICP increased monotonically with each infusion step, reaching severe elevations during the terminal stage (3 ml/min). ABP initially increased in parallel with ICP, reflecting systemic compensatory mechanisms, while rCBF also increased during the early infusion stages (0.05-0.3 mL/min) in association with ABP elevation. However, during the 1.0 mL/min stage, rCBF showed reduced correlation with ABP and an inverse correlation with ICP, indicating a gradual loss of ABP-mediated compensation. During the terminal stage, ICP and rCBF inversely mirrored each other, indicating loss of CA.

Zoom-in waveform analysis during baseline reveals clear cardiac pulsatility in rCBF that closely tracked simultaneous ABP and ICP waveforms (**Fig. 3d**), highlighting the ability of TR-LSCI to resolve pulsatile microvascular flow dynamics at high temporal resolution. A small phase offset was observed among pulsatile waveforms across three modalities, attributable to differences in measurement location (cerebral rCBF and ICP vs. peripheral femoral ABP), resulting in physiological pulse transit delays. Additional contributions include modality-specific sensor dynamics and minor timing offsets during manual alignment of acquisition markers. However, these phase differences do not affect individual waveform morphology or HR extraction across modalities.

**Fig. 4** shows another representative adult rat (Rat #2), which demonstrates similar physiological responses to the intracranial infusion. In this animal, saline was infused sequentially at 0.05 mL/min (∼20 min), 0.1, 0.2, and 0.3 mL/min (each ∼5 min), followed by 1.0 mL/min (∼3 min). This infusion protocol was similar to that used for Rat #1, with stepwise increases in infusion rate; however, the maximum infusion rate reached differed across animals due to inter-animal variability in physiological tolerance to intracranial loading.

Baseline cortical BFI within the defined ROI (**Fig. 4a**) exhibited a relatively uniform spatial distribution (**Fig. 4b**). With increasing infusion rates, cortical BFI showed progressive redistribution and augmentation during low-to-moderate infusion stages, followed by attenuation during the 1.0 mL/min stage. For this rat, marked physiological instability emerged during the 1.0 mL/min stage, and the experiment was therefore terminated without proceeding to a higher infusion rate. The synchronized ICP, ABP, and rCBF time courses (**Fig. 4c**) and baseline zoom-in cardiac waveforms (**Fig. 4d**) similarly illustrate preserved pulsatile coupling during baseline and early infusion, with breakdown of flow regulation as ICP increased. Although inter-animal variability in absolute signal magnitude and spatial pattern was observed, the overall trends in rCBF evolution and pressure-flow interactions were consistent across all four rats.

### 3.2 rCBF, ABP, and ICP Dynamics in A Neonatal Piglet During Intraventricular Infusion

**Fig. 5** presents the results from a neonatal piglet subjected to an intermittent saline infusion protocol, demonstrating the translational applicability of the mobile TR-LSCI system in a larger, immature brain. **Fig. 5a** shows the raw TR-LSCI intensity image with the selected cortical ROI for subsequent time-course data analysis.

**Fig. 5.**
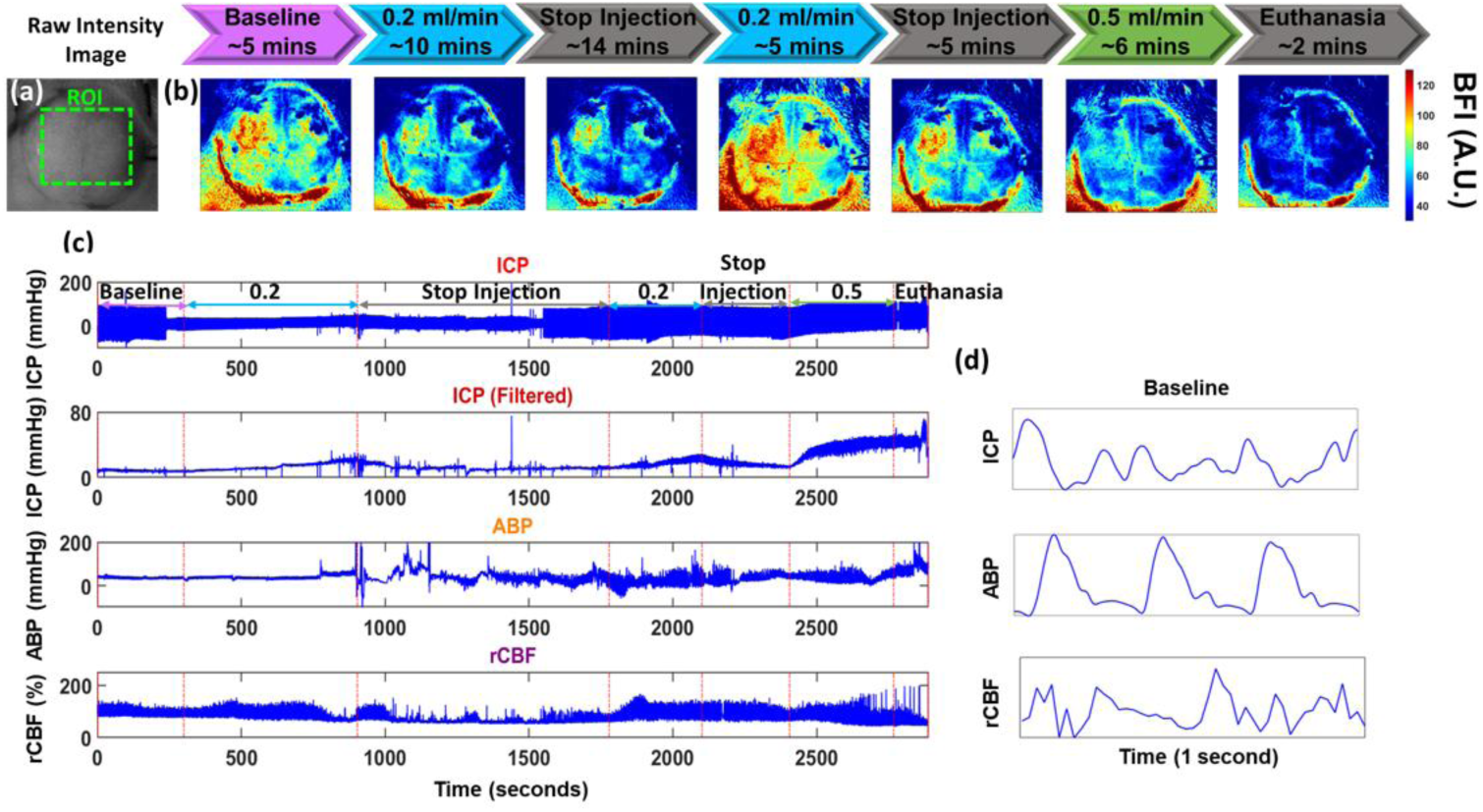
Piglet results demonstrating two-dimensional BFI maps and synchronized multimodal physiological responses during graded intracranial infusion. **(a)** Raw TR-LSCI intensity image showing the cortical ROI used for time-course data analysis. **(b)** Two-dimensional BFI maps acquired over the intact skull at baseline and during repeated infusion and pause cycles: 0.2 ml/min for 10 min, stop (14 min), 0.2 ml/min for 5 min, stop (5 min), 0.5 ml/min for 6 min, followed by terminal phase. All BFI maps were derived from Gate #40 and represented averaging over the final 20 s of each infusion segment. **(c)** Time courses of ICP, ABP, and rCBF, showing dynamic cerebrovascular responses to intermittent infusion. Vertical dashed lines indicate infusion transitions. The upper ICP trace shows the raw signal, while the second ICP trace reflects the filtered signal after 60 Hz notch and 25 Hz low-pass filtering to remove instrumentation-related noise. Sampling rates: ICP and ABP at 1000 Hz; rCBF at 46 Hz with three depth gates and 8-bit acquisition. **(d)** Zoom-in waveform signals for ICP, ABP, and rCBF at the baseline period of 1 sec.

At baseline, cortical BFI within the defined ROI **(Fig. 5a)** derived from the deeper gate (Gate #40) showed a relatively uniform spatial distribution **(Fig. 5b).** During the initial infusion stage (0.2 ml/min for ∼10 min), ICP increased steadily until 30 mmHg and was accompanied by a moderate elevation in BFI with clear spatial redistribution of cortical flow. Following cessation of infusion (∼14 min), both ICP and BFI partially recovered toward baseline levels. A second infusion at the same rate (0.2 ml/min for ∼5 min) elicited a similar but more pronounced, quicker elevation of ICP to 30 mmHg, again followed by partial recovery during a shorter pause (∼5 min). The final infusion at 0.5 ml/min (∼6 min) induced a rapid rise in ICP, accompanied by marked disruption of pressure-flow coupling and widespread suppression of cortical BFI preceding terminal stage.

The synchronized physiological time-course data quantified these dynamics (**Fig. 5c**). Raw ICP recordings exhibited an instrument electrical power noise at 60 Hz, which was removed using a 60-Hz notch filter followed by a 25 Hz low-pass filter; both raw and filtered traces are shown for comparison. Across infusion cycles, ICP closely tracked injection epochs, while ABP displayed complex compensatory behavior characterized by transient fluctuations during both infusion and pause periods. rCBF time-course data demonstrated a multiphasic response, with moderate augmentation during early infusion, partial recovery during pauses, and eventual collapse during the final high-rate infusion. Zoom-in pulsatile waveforms revealed preserved cardiac pulsatility during baseline measurements (**Fig. 5d**).

When analysis was restricted to injection periods, the piglet exhibited greater spatial heterogeneity in BFI, broader cortical flow redistribution, and more dynamic ICP and ABP fluctuations compared with the rats. These differences primarily reflect the necessity of a large-animal model to more clearly reveal the dynamic coupling among ICP, ABP, and CBF under clinically relevant ICP manipulations, rather than species-specific recovery effects. Despite differences in species, age, infusion strategy, and tolerated ICP range, both models demonstrated reproducible and structured rCBF responses to progressive intracranial loading.

### 3.3 Cardiac Pulsatility and Heart-Rate Consistency Across rCBF, ICP, and ABP

HR was independently extracted from ICP, ABP, and rCBF signals during baseline using a sliding-window FFT. HR estimates from all three modalities showed strong agreement (**Table 1**), indicating that TR-LSCI derived rCBF preserves cardiac pulsatility and yields physiologically consistent HR estimates compared with standard pressure measurements.

**Table 1:**
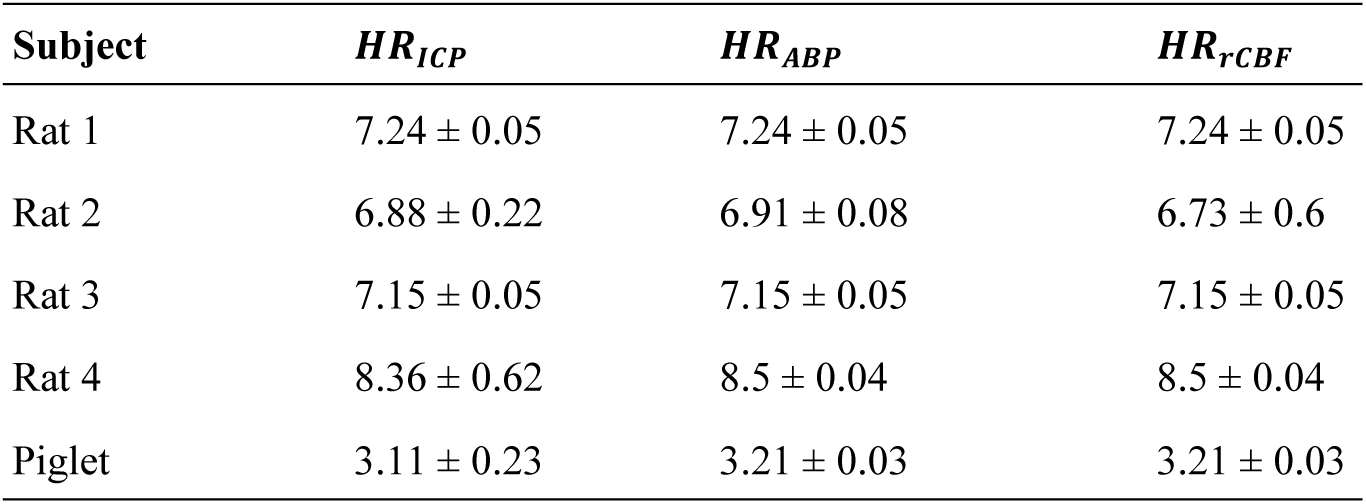
Baseline heart rates (mean ± standard deviation) derived from ICP, ABP, and rCBF.

The across-rat TOST equivalence testing further quantified this agreement (**Table 2**). For all three pairwise comparisons (ABP vs rCBF, ICP vs rCBF, and ABP vs ICP), mean differences were small, and the corresponding 90% confidence intervals lay entirely within the predefined equivalence bounds of ±0.25 Hz.

**Table 2:** Across-rat equivalence testing of HR estimates (TOST)

| Comparison | 90% CI (Hz) | p ($> -0.25$ Hz) | p ($< +0.25$ Hz) | Equivalence |
| --- | --- | --- | --- | --- |
| ABP vs rCBF | [-0.061, 0.151] | 0.0036 | 0.0099 | Yes |
| ICP vs rCBF | [-0.137, 0.142] | 0.0118 | 0.0125 | Yes |
| ABP vs ICP | [-0.036, 0.121] | 0.0015 | 0.0042 | Yes |

Within-subject (Across-time-point) TOST analyses demonstrated consistent equivalence across baseline time points. For all rats, all pairwise HR comparisons were statistically equivalent across thousands of baseline time points, with narrow confidence intervals and highly significant one-sided tests (p < 0.001). Similarly, all pairwise HR comparisons in the piglet dataset were statistically equivalent across baseline time points (p < 0.001).

Collectively, these results demonstrate that noninvasive TR-LSCI derived rCBF reliably preserves cardiac pulsatility and yields HR estimates that are statistically and physiologically equivalent to those obtained from invasive ICP and ABP measurements, both at the group level and over time within individual subjects.

### 3.4 Multimodal Monitoring Reveals Distinct CA Phases During Graded ICP Loading

**Fig. 6** summarizes the dynamics of ΔBFI, ΔABP, and ΔICP across all animal experiments, revealing similar pressure-flow regulation patterns under progressive intracranial loading. These dynamics were categorized into three primary CA phases, with an additional transient period (T-period).

**Fig. 6.**
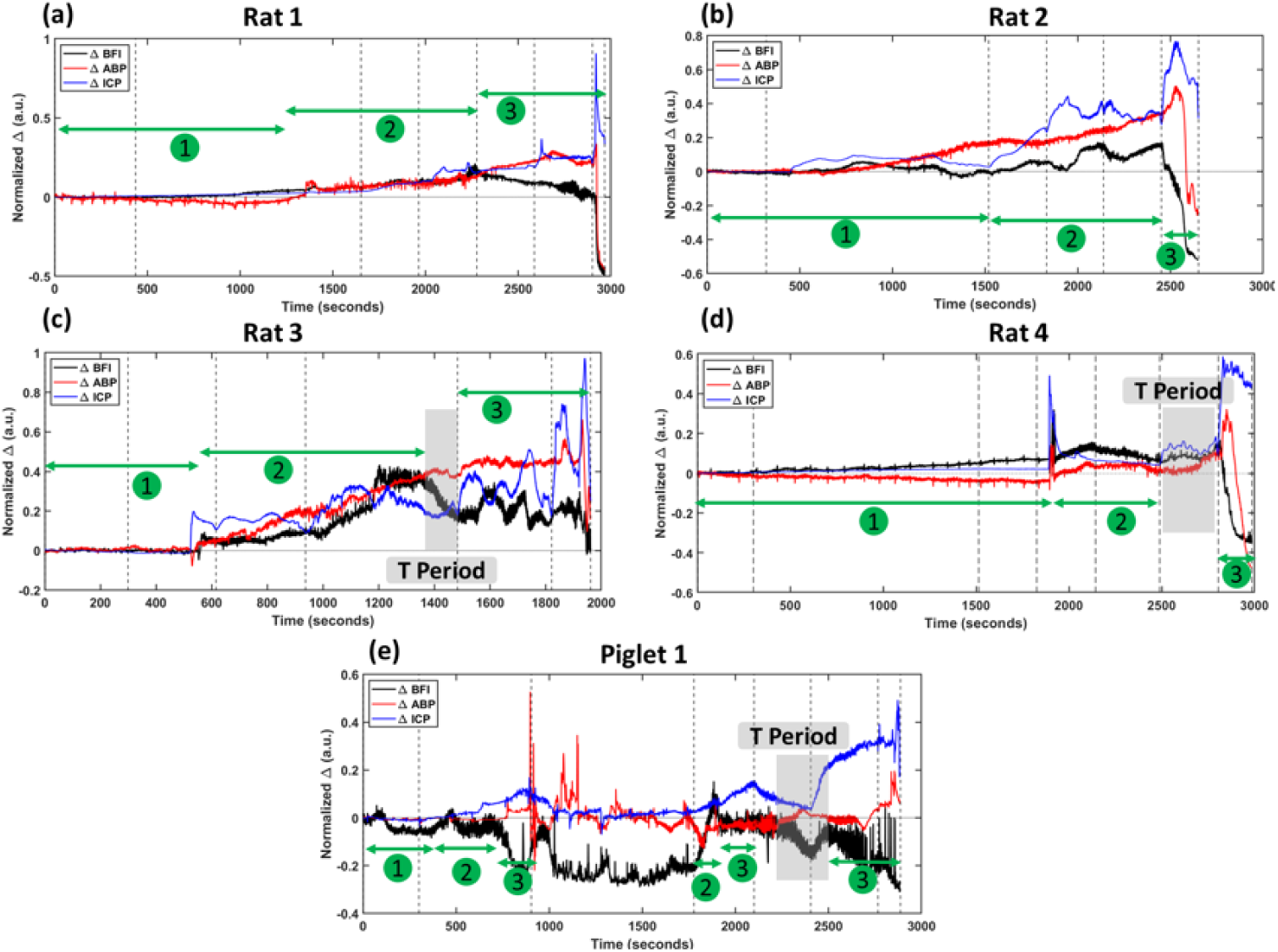
Phase-dependent pressure-flow coupling revealed by interaction analysis of ΔBFI, ΔABP, and ΔICP. (a-d) ΔBFI (black), ΔABP (red), and ΔICP (blue) for four adult rats and **(e)** the corresponding analysis for the neonatal piglet, during graded intracranial pressure loading. Vertical dashed lines indicate infusion step transitions. Signals were min-max normalized and baseline-subtracted to enable direct qualitative comparison across modalities. Green arrows labeled **①-③** indicate reproducible three CA phases: **①** Phase 1 (intact CA), in which ΔBFI remains buffered despite gradual changes in ΔABP and ΔICP; **②** Phase 2 (impaired CA), characterized by ABP-coupled CBF augmentation, where ΔBFI increases in parallel with ΔABP despite rising ΔICP; and **③** Phase 3 (loss of CA), marked by ICP-constrained CBF suppression with ΔBFI plateauing or declining as ΔICP rises steeply. Gray shaded regions denote transient period (T), during which ΔBFI cannot be clearly attributed to either ABP or ICP alone. The timing and duration of these phases varied across animals.

During Phase 1, ΔBFI remained relatively stable despite gradual increases in both ΔABP and ΔICP. In this interval, CBF responses were buffered against pressure fluctuations, and no strong coupling to either pressure signal was observed, consistent with preserved CA. Physiologically, this phase reflects intact autoregulatory vasoreactivity, where cerebral vessels adjust resistance to maintain stable flow despite changing perfusion pressures.

As intracranial loading progressed, CA entered Phase 2 with the ABP-coupled CBF augmentation. In this phase, ΔBFI increased in parallel with elevated ΔABP while ΔICP continued to rise, indicating enhanced coupling between ABP and CBF. Importantly, CBF augmentation occurred despite concurrent ICP elevation, suggesting that ABP compensation remained effective and that ICP had not yet imposed a dominant mechanical constraint on CBF. This Phase 2 reflects impaired CA, in which autoregulatory buffering is weakened and CBF becomes increasingly dependent on ABP.

With further ICP elevation, CA entered Phase 3 with the ICP-constrained CBF suppression. ΔICP rose steeply while ΔBFI plateaued or declined, even as ΔABP remained elevated. This pattern reflects the limitation of CA, consistent with exhaustion of autoregulatory reserve and increasing intracranial constraint on microvascular flow. In this Phase 3, CA is lost, and elevated ICP mechanically restricts CBF regardless of systemic pressure compensation.

In addition to these three major phases, the gray-shaded intervals highlight transient periods between Phase 2 and Phase 3 with indeterminate pressure-flow coupling rather than a distinct autoregulatory phase. During these periods, ΔBFI exhibited rapid fluctuations or mixed responses that could not be clearly attributed to either ABP or ICP alone. These transition intervals likely reflect dynamic vascular adjustments occurring during rapid physiological transitions, such as changes in infusion rate or abrupt shifts in intracranial compliance and were therefore classified as transient periods rather than distinct regulatory phases.

Notably, the timing and prominence of these phases varied across subjects. For example, Rat 3, which did not undergo the prolonged low-rate (0.05 ml/min) infusion, exhibited an accelerated loss of autoregulatory ability (Phase 1), transitioning more rapidly from Phase 1 to Phase 3. Similarly, in the piglet experiment, autoregulatory ability was markedly attenuated during the second infusion cycle, with CBF entering Phase 3 earlier than in the initial injection and a reduced duration of Phase 2. This earlier CA breakdown in the piglet likely reflects the combined effects of repeated intracranial loading, larger brain volume, and immature cerebrovascular control.

Together, these results demonstrate that interaction analysis of ΔBFI, ΔABP, and ΔICP provides a powerful framework for visualizing phase-dependent cerebrovascular regulation, capturing both reproducible physiological patterns and subject-specific variability across experimental protocols.

## 4 Discussion and Conclusions

Reliable assessment of CA requires continuous evaluation of how CBF responds to evolving intracranial and arterial pressure conditions. In neurocritical care, the inability to track autoregulatory state in real time limits the development of targeted, patient-specific interventions and contributes to secondary brain injury during periods of intracranial instability. Within this context, there is a clear need for methods that can resolve cerebrovascular dynamics continuously fast enough, at the level of the microvasculature with high spatial resolution, and under physiologically relevant perturbations [21, 53].

Within this framework, the coupled behavior of CBF, ABP, and ICP can be viewed as a physiological reference standard for interrogating cerebrovascular regulation. The central objective of the present study was therefore to establish this coupled behavior experimentally by combining scalable, noncontact, wide-field, depth-sensitive TR-LSCI imaging of microvascular CBF with high temporal and spatial resolution and synchronized invasive measurements of ICP and ABP. This integration enables interpretation of rCBF dynamics explicitly in the context of concurrent pressure perturbations, rather than inferring cerebrovascular state from pairwise correlations or indirect surrogates [54–56].

As illustrated in **Fig. 1**, TR-LSCI enables wide-field imaging of microvascular CBF, while concurrent ICP and ABP measurements provide independent pressure readouts, allowing direct interrogation of triadic pressure-flow interactions beyond conventional pairwise autoregulation indices [25–29, 31–40]. To support both controlled mechanistic studies and translationally relevant experiments, we developed complementary benchtop and mobile TR-LSCI systems for adult rat and neonatal piglet models, respectively, building on our prior platform development [49, 50]. While the benchtop configuration provided a stable environment for systematic physiological interrogation, translation to a mobile, fiber-coupled system was essential for extending TR-LSCI to larger animals and for imaging under clinically relevant conditions.

The integrated signal-processing and analysis framework (**Fig. 2**) was essential for extracting physiologically meaningful rCBF dynamics and enabling reliable multimodal comparison with ICP and ABP. Depth-gated speckle processing and ROI-based rCBF extraction improved SNR while preserving both slow hemodynamic trends and cardiac pulsatility, which are critical for characterizing cerebrovascular regulation. Synchronization and preprocessing of pressure signals ensured temporal correspondence across modalities, enabling direct assessment of flow-pressure interactions. Together, this unified pipeline provided a robust foundation for validating rCBF against invasive measurements and for revealing phase-dependent transitions in CA during graded intracranial loading.

Using this integrated platform, we first characterized cerebrovascular responses in adult rats subjected to continuous graded intraventricular infusion (**Figs. 3-4**). These experiments demonstrate that TR-LSCI can resolve both spatially heterogeneous microvascular flow patterns and high-frequency pulsatile dynamics during progressive ICP elevation. Early infusion stages were marked by preserved or only mildly impaired autoregulation, whereas higher infusion rates revealed increasing ABP-CBF coupling and eventual flow suppression, consistent with progressive loss of autoregulatory reserve.

The neonatal piglet experiment (**Fig. 5**) extended these observations to a larger brain under intermittent loading conditions that more closely resemble clinical scenarios. Compared with rats, the piglet exhibited greater spatial heterogeneity, more pronounced transient ABP responses, and earlier attenuation of autoregulatory behavior during repeated ICP challenges. These differences likely reflect developmental factors, larger brain volume, thicker skull, and altered intracranial compliance, underscoring the importance of large-animal models for translational cerebrovascular research. Despite these distinctions, the fundamental pressure-flow patterns were conserved across models, supporting the physiological generality of the observed autoregulatory behavior.

The physiological validity of TR-LSCI derived rCBF was independently verified through cardiac pulsatility analysis. Heart rate extracted from rCBF closely matched HR derived from invasive ICP and ABP measurements in both rats and the piglet, with statistical equivalence confirmed across subjects and within baseline recordings (**Tables 1-2**). This agreement demonstrates that TR-LSCI preserves cardiac-frequency microvascular flow dynamics and supports interpretation of the observed pressure-flow relationships as genuine cerebrovascular regulation rather than measurement artifact.

The central physiological finding of this study is that cerebrovascular regulation evolves through three reproducible CA phases, sometimes separated by transient periods (**Fig. 6**). Phase 1 is characterized by stable ΔBFI despite rising ΔICP and ΔABP, consistent with intact autoregulatory vasoreactivity. Phase 2 exhibits ABP-coupled CBF augmentation (ΔBFI), reflecting impaired but still compensatory autoregulation in which systemic pressure elevation (ΔABP) offsets intracranial pressure loading (ΔICP). Phase 3 is marked by ICP-constrained CBF suppression, indicating exhaustion of autoregulatory reserve and mechanical limitation of microvascular perfusion.

Importantly, these phases reflect distinct physiological regimes of cerebrovascular control rather than arbitrary segmentation. The observed ABP elevation accompanying ICP increases is consistent with emerging concepts such as the brain baroreflex, in which autonomic cardiovascular responses act to counter intracranial challenges and preserve cerebral perfusion [57, 58]. Our data provide experimental support for this framework by directly demonstrating how ABP-driven compensation can transiently sustain or augment CBF before ICP constraints dominate. The transient periods further emphasize that CA is dynamic rather than binary, with rapid fluctuations and mixed coupling during changes in infusion rate or intracranial compliance.

An important translational insight from this study is that longitudinal rCBF dynamics alone contain substantial information about CA status, even in the absence of direct pressure measurements. Across both rats and the piglet, consistent phase-specific rCBF patterns were observed, including relative flow stability during preserved autoregulation (Phase 1), progressive augmentation during compensatory responses (Phase 2), and abrupt or sustained suppression during autoregulatory failure (Phase 3). These observations suggest that continuous CBF monitoring may enable qualitative inference of autoregulatory status and impending deterioration when interpreted longitudinally and in the context of known interventions.

At the same time, the findings underscore the limitations of CBF-only inference. Similar rCBF trajectories can arise from different combinations of ICP, ABP, and vascular compliance, particularly during transient or unstable periods. Accordingly, while rCBF dynamics alone provide valuable insight, concurrent pressure measurements, when clinically available, remain important for mechanistic interpretation and validation in complex or rapidly evolving pathological states.

The present study has some limitations related to experimental scope and data analysis. The large-animal component of this study was limited to a single neonatal piglet, which restricts statistical generalization across subjects and developmental stages. While the piglet experiment was intended to demonstrate feasibility and translational relevance rather than population-level inference, future studies with larger cohorts will be required to assess inter-subject variability, maturation-dependent effects, and disease-specific alterations in CA. In addition, although phase-dependent transitions in CA were consistently observed, inter-signal phase relationships among rCBF, ABP, and ICP waveforms were evaluated qualitatively rather than quantitatively. Quantitative analysis will require absolute CBF measurements, which will be addressed in future studies, for example by using multi-exposure approaches to quantify absolute BFI [59–62].

Several other directions for future work naturally follow from this study. Data-driven estimation of ICP from optical rCBF signals is a promising direction, building on prior diffuse optics studies using pulsatile dynamics and cardiac timing [38, 40, 63]. Machine learning models applied to the spatiotemporal features of TR-LSCI rCBF could enable noninvasive ICP estimation by learning complex pressure-flow relationships that are difficult to model analytically. Explicit synchronization strategies (e.g., EKG-based triggering or shared hardware timing) will enable pulse-and phase-resolved analyses by ensuring precise temporal alignment between rCBF waveforms and cardiac and pressure dynamics [38, 40]. Additional extensions include multi-wavelength TR-LSCI for simultaneous assessment of CBF, cerebral oxygenation, and cerebral metabolic coupling, as well as next-generation SPAD cameras such as SPAD Alpha with higher pixel density (1024 × 1024) and fill factors to improve spatial resolution and depth sensitivity. Together, these advances will support the development of a fully noninvasive optical imaging platform for continuous cerebrovascular monitoring in neonates and other vulnerable patient populations.

In conclusion, this study demonstrates that TR-LSCI enables noncontact, depth-resolved, high-speed, high-resolution imaging of CBF and, when combined with concurrent ICP and ABP measurements, provides a comprehensive framework for investigating dynamic CA. Through integrated system design, unified analysis pipeline, and multimodal physiological measurements, we show that CA progresses through reproducible, phase-dependent transitions from preserved autoregulation to ABP-driven compensation, and ultimately to ICP-constrained CBF suppression. Importantly, CBF dynamics alone exhibit distinct signatures corresponding to these phases, indicating that continuous CBF monitoring can yield meaningful insight into CA even in the absence of invasive pressure measurements. While simultaneous ICP and ABP monitoring remains important for definitive mechanistic interpretation, mobile TR-LSCI offers a promising noninvasive approach for tracking autoregulatory integrity and identifying early deterioration in both preclinical and translational settings. Together, these findings establish TR-LSCI as a powerful platform for dynamic, physiology-informed neurovascular monitoring and lay the foundation for future optical strategies aimed at bedside CA assessment.

## Acknowledgments

We acknowledge partial financial support from the National Institutes of Health (NIH) R01-EB028792, R01-HD101508, R21-HD091118, R21-NS114771, R41-NS122722, R42-MH135825, R56-NS117587 (G.Y.), and the Halcomb Fellowship in Medicine and Engineering at the University of Kentucky (F.F.). This work was also supported, in part, by the Swiss National Science Foundation (grants 20QT21_187716 Qu3D “Quantum 3D Imaging at high speed and high resolution” and 200021_166289).

## Disclosure

G.Y. serves as an advisor for Bioptics Technology, which is developing products related to the research being reported. G.Y.’s family has equity ownership in Bioptics Technolgy. L.C. (Lei Chen) serves as a consultant to the Bioptics Technology. The terms of this arrangement have been reviewed and approved by the University of Kentucky.

Edoardo Carbon is co-founder of Novoviz, and Edoardo Charbon and Claudio Bruschini are co-founders of Pi Imaging Technology. Neither company has been involved with the work nor the paper drafting.

## Data Availability

Data underlying the results presented in this paper are not publicly available at this time but may be obtained from the authors upon reasonable request.

## Notes

### Competing Interest Statement

The authors have declared no competing interest.

